# Integrating cold hardiness and deacclimation resistance demonstrates a conserved response to chilling accumulation in grapevines

**DOI:** 10.1101/2024.09.28.615590

**Authors:** Jason P Londo, Al P Kovaleski

## Abstract

To survive the harsh conditions of winter, woody perennial species such as grapevine have adapted to use environmental cues to trigger physiological changes to induce dormancy, acquire cold hardiness, and measure the length of winter to properly time spring budbreak. Human induced climate change disrupts these cues by prolonging warm temperatures in fall, reducing the depth and consistency of midwinter, and triggering early budbreak through false spring events. We evaluated variation in dormant bud cold hardiness and chilling hour requirements of 31 different grapevine varieties over 3 years. Differential thermal analysis was used to track changes in cold hardiness and deacclimation resistance was assessed throughout the season to track dormancy progression. Results demonstrate wide variation in maximum deacclimation rate (1.03 – 2.87 °C/day) among varieties under forcing conditions. Absolute maximum rates of deacclimation show signatures of species-level responses to forcing temperatures. When integrated with variation in cold hardiness, these rates revealed a relationship between winter cold hardiness, changes in deacclimation rate and budbreak phenology. Standardizing rates among varieties as deacclimation potential demonstrated a conserved response to chilling exposure among varieties that alters our interpretation of the concept of high and low chill varieties and chilling requirement in grapevine.

## Introduction

The impact of changing climate on perennial fruit crops is complex and while heat and drought are major challenges for sustainable crop production, studies demonstrate that climate warming is greater during the winter season (Marvel *et al*., 2023). Milder winters may offer a potential benefit for cool temperate fruit production, reducing the chances of midwinter freeze damage. However, warming climate increases the frequency of midwinter warming events, termed false springs (Rochette *et al*., 2004; Gu *et al*., 2008; Augspurger, 2013), which contribute to substantial freezing damage as perennial crops rely on cold temperatures to maintain dormancy and freeze resistance (Gu *et al*., 2008; Vitasse *et al*., 2014; Vyse *et al*., 2019). Additionally, winter climate warming contributing to advances in spring phenology, increased risk of early spring frost damage (Bigler and Bugmann, 2018; De Rosa *et al*., 2021; Lamichhane, 2021).

European grapevine (*Vitis vinifera* L.) is one of the most economically important fruit crops in the world with production on an estimated 7.5 million hectares, resulting in 85 million metric tons of fruit (‘FAOSTAT’; Grassi and De Lorenzis, 2021). *Vitis vinifera* was domesticated from its wild ancestor *V. vinifera* spp. *sylvestris* roughly 8000 years ago in the Near East and Caucasus region (Grassi and De Lorenzis, 2021). Preferentially adapted to its climate of origin – the hot dry summers and cool mild winters of a Mediterranean climate – the European grapevine has been spread by humans around the globe to result in major grape production regions on every continent, absent Antarctica. Given the crop’s relevance, grapevines are often the subject of climate change concern and several studies have focused on the potential loss of viticultural regions as global temperatures rise (Hannah *et al*., 2013; Mozell and Thach, 2014; Morales-Castilla *et al*., 2020; Keller, 2023), though other studies document the adaptability of grapevine (van Leeuwen *et al*., 2013; Keller, 2023). In addition to *V. vinifera* and its wild ancestor, the genus *Vitis* includes 80 wild species split into two large clades with diversification in Asia and North America (Govaerts, 2024). While *V. vinifera* is prized for its high-quality fruit, the species lacks much of the diversity in disease resistance and climate adaptability that can be found within wild grapevine. As a result, grapevine breeding for the development of new cultivars often draws on wild grapevine germplasm to produce regionally adapted cultivars.

Grapevine breeding has as long history in North America (Reynolds and Reisch, 2015) and resistance to winter temperatures has been a target of grape breeding in cool climates like the Eastern United States. Historically, midwinter temperatures in these regions exceed −25°C, a temperature that has been shown to cause freezing damage in the dormant buds of many popular *V. vinifera* cultivars (Mills *et al*., 2006; Ferguson *et al*., 2011; Londo and Kovaleski, 2017). Wild grapevine species native to North America and which have northern distributions, including *V. riparia*, *V. labrusca*, and *V. aestivalis,* are adapted to deep midwinter freezing temperatures and can survive winter temperatures as low as −30 to −35°C (Londo and Kovaleski, 2017). Breeding with these wild species increases the midwinter hardiness of the resulting hybrid cultivars and many modern North American hybrid cultivars include these wild species in their pedigree (Reynolds and Reisch, 2015). However, these northern wild grape species, and many of the hybrid cultivars derived from them, tend to lose cold hardiness quickly in late winter and break bud early in the spring, potentially exposing them to greater frost risk (Londo and Kovaleski, 2017; Kovaleski *et al*., 2018; De Rosa *et al*., 2021). While long a target of breeding and research, the genetic architecture of cold hardiness in grapevine remains unclear.

Grapevines and other perennial species undergo specific physiological changes to induce and maintain dormancy throughout winter while developing cold hardiness, enabling them to survive temperatures far below freezing. Dormancy, defined as the cessation of visible growth, occurs as three broadly different categories: paradormancy, endodormancy, and ecodormancy (Lang *et al*., 1987). Paradormancy is the suppression of growth due to hormonal or other physiological factors during the growing season, such as the suppression of lateral buds due to apical dominance. Grapevine buds transition from paradormancy to endodormancy at the end of the growing season in response to decreasing photoperiod and daily temperatures (Fennell, 2004; Garris *et al*., 2009). Growth and development during endodormancy are restricted due to currently unknown endogenous mechanisms, but these mechanisms protect the bud from responding to fluctuating environmental conditions (Horvath *et al*., 2003; Anderson *et al*., 2010).

In parallel with endodormancy induction, grapevine buds develop the ability to survive freezing temperatures through a mechanism called supercooling (Andrews *et al*., 1984; Wisniewski *et al*., 2003). This process involves controlled dehydration and physical isolation of the bud primordia from the vascular system of the cane, enabling the suppression of freezing of intracellular water (Jones *et al*., 2000; Rubio *et al*., 2016). In early winter, dormant buds progressively gain greater freeze resistance in response to declining temperatures, a process known as acclimation (Wisniewski *et al*., 2003, 2014). Cold hardiness is maintained through midwinter through currently unknown mechanisms. To successfully transit winter and break bud in the spring, grapevines must transition from the growth repression of endodormancy, to ecodormancy, where growth repression is maintained by the unsuitable (low temperature) environmental conditions of late winter (Lang *et al*., 1987; Campoy *et al*., 2011). Ecodormancy thus represents a precarious stage as it relates to warming winter conditions. If ecodormant buds experience warm temperatures, they begin to lose cold hardiness, known as deacclimation, in a temperature-dependent manner (Kalberer *et al*., 2006; Pagter and Arora, 2013; Kovaleski *et al*., 2018). Understanding the timing of the endodormancy to ecodormancy transition is critical to understanding how climate change will impact the cold hardiness and resilience of grapevines during the winter season.

The molecular mechanisms that drive the transition between endodormancy and ecodormancy are unknown. However, it is well established that exposure to low, non-freezing temperatures, called chilling, plays a critical role in this timing. The quantity of chilling exposure is termed the chilling requirement (CR) and a crop or cultivar’s CR is typically determined by observing the time needed for dormant plant tissues to reach budbreak under growth-permissive conditions. Buds are considered ecodormant and the CR met when 50% of total budbreak occurs within a specific temporal threshold (Lavee and May, 1997; Dennis, 2003; Pellegrino *et al*., 2020). Chilling requirements (CR) are crucial for determining cultivar suitability, as cultivating plants with high chill needs in low chill climates can result in poor budbreak synchronization and disrupted phenology during the growing season (Campoy *et al*., 2011; Luedeling *et al*., 2013; Pertille *et al*., 2021; Drogoudi *et al*., 2023). Conversely, using low chill cultivars in high chill environments often leads to early budbreak and increased frost risk (Atkinson *et al*., 2013; Fernandez *et al*., 2023).

Various chilling models can be used to estimate winter chilling accumulation, differing by crop type and region. These models range from simple cumulative measures of hours spent within temperature thresholds (e.g., Chilling Units (Dokoozlian, 1999)) to complex multistep methods that combine temperature thresholds and temperature variation to model the response (e.g., Dynamic Portions Model (Erez *et al*., 1990)). In grapevines, chilling requirements have been estimated for cultivated (Dokoozlian, 1999; Yilmaz *et al*., 2020; Anzanello *et al*., 2021; Rahemi *et al*., 2021) and wild species (Londo and Johnson, 2014). Climate change is expected to impact chilling accumulation rates and quantities. For example, warmer winter conditions are predicted to alter chilling exposure across regions, with warm areas likely experiencing reduced chilling and cold regions potentially gaining chilling exposure (Luedeling, 2012; Atkinson *et al*., 2013; Chamberlain and Wolkovich, 2021; Delgado *et al*., 2021; Fraga and Santos, 2021). Studies show that while some models effectively estimate regional chilling requirements, few accurately model chilling across diverse production regions (Luedeling and Brown, 2011; Luedeling, 2012). This limitation highlights a fundamental gap in our understanding of chilling accumulation mechanisms and the physiological impact of chilling on dormancy.

Winter physiology is a complex interaction between genotype and environment, influencing the traits of maximum cold hardiness, dormancy, and budbreak phenology. Recently, studies in grapevines (Kovaleski *et al*., 2018; Kovaleski and Londo, 2019; North *et al*., 2022), and more broadly among 15 woody perennial species (Kovaleski, 2022) demonstrated connections between dormant bud cold hardiness, chilling accumulation, deacclimation rate, and budbreak phenology. These interactions highlight several unanswered questions. How different are the chilling requirements of *V. vinifera* from hybrid cultivars? Are *V. vinifera* or hybrid cultivars at higher risk of climate change induced winter warming? How is cold hardiness, dormancy status, and budbreak phenology linked in grapevine? To answer these questions, we characterized the cold hardiness of 31 grapevine cultivars at weekly time points across three winters using differential thermal analysis. In tandem, we measured the resistance to deacclimation under forcing temperatures and budbreak phenology relative to chilling accumulation. Combining these traits, we capture the subtle changes in dormancy state that occur as buds transition from endodormancy to ecodormancy and test the potential integration of cold hardiness and deacclimation rates to predict budbreak phenology.

## Materials and Methods

All data processing and analysis in this study was conducted using R Statistical Software (v4.2.1; R Core Team 2022) with the following packages: tidyverse, ggplot2, chillR, lsmeans, dormancyR, drc, lubridate, broom, and patchwork.

### Weather data and chilling models

Hourly temperature data was retrieved from the Network for Environment and Weather Applications (NEWA; http://newa.cornell.edu/) website for the Geneva, New York, United States station (Bejo) for all three years of the study (2018-2021). Estimates of chilling hours in the three winters were evaluated for various chill models. Chill units for Utah and Dynamic portion (DP) models were computed using the chillR package (‘chillR Package: Statistical Methods for Phenology Analysis in Temperate Fruit Trees’, 2022) while the North Carolina (NC) and Chilling units (CU) were computed using the dormancyR package (Fernandez *et al*., 2020). In all cases, the winter season was set to start on 1 September and run through 30 April.

### Plant Sampling

This study was conducted over three consecutive winter seasons: 20 September 2018-16 April 2019 (Year 1), 5 September 2019-2 April 2020 (Year 2), and 2 October 2020-31 March 2021 (Year 3). Thirty-one cultivars of grapevine were sampled from local commercial and experimental vineyard locations (**Table S1**) at approximately weekly intervals in each of the three years to evaluate cold hardiness in response to natural field conditions, deacclimation resistance, and budbreak phenology under controlled temperature conditions. Cultivars included both red (R) and white (W) fruited European *Vitis vinifera* cultivars (hereafter, *vinifera*); Cabernet Franc (R), Cabernet Sauvignon (R), Chardonnay (W), Gewurztraminer (W), Gruner Veltliner (W), Lemberger (R), Merlot (R), Pinot gris (W), Pinot noir (R), Riesling (W), Sangiovese (R), Saperavi (R), Sauvignon blanc (W), Syrah (W), and Tocai Friulano (W), and hybrid cultivars (hereafter, *hybrid*) with various wild species genetic backgrounds; Aromella (W), Cayuga White (W), Chambourcin (R), Chancellor (R), Concord (R), Corot noir (R), La Crescent (W), Marechal Foch (R), Marquette (R), Niagara (W), Noiret (R), St. Croix (R), Traminette (W), Valvin Muscat (W), Vidal (W), and Vignoles (W). These cultivars represent a subsample of the diversity of cultivars grown in the Northeastern United States and were chosen in part to also represent diversity in optimum climate niche. For the *vinifera* cultivars, we included both “cool” and “warm” climate varieties (Jones *et al*., 2012) and for *hybrid* cultivars, we used cultivars that represent both French American hybrids and New American hybrids, as well as hybrids derived from wild grapevine species that have Northern (*V. labrusca* or *V. riparia*), or Southern (*V. rupestris* or *V. lincecumii*) distributions.

Distributions of wild species were obtained from county level data found at the Biota of North America Program (https://bonap.net/napa). Pedigree data for hybrid cultivars was retrieved from the Vitis International Variety Catelog (VIVC, https://www.vivc.de/). All cultivar information is included in **Table S1.**

Dormant cane cuttings with nodes 3-20 (depending on cultivar and pruning method) were collected from the field and transported to the bud physiology lab at Cornell AgriTech for processing. During each collection time point in 2018-2019, 40 nodes of each cultivar were collected from the field by randomly pruning from at least 4 different individual vines at each collection. In 2019-2020 and 2020-2021, sampling was increased to 50 nodes of each cultivar at each collection. Upon returning from field collections, dormant cane/node material were promptly pruned into single node cuttings, randomized, and distributed into subsamples for experiments to determine an initial cold hardiness assessment (n = 5; 2018-2019, n = 10; 2019-2020 and 2020-2021), deacclimation resistance (n =25), and budbreak phenology (n =10).

### Assessment of cold hardiness using differential thermal analysis

Cold hardiness response curves for each year and each cultivar were produced using the mean LTE value for the weekly collection (T0) timepoints. For determining dormant bud cold hardiness levels, grapevine supercooling ability was assessed following standard practices for differential thermal analysis (DTA) (Mills *et al*., 2006; Ferguson *et al*., 2011; Londo and Kovaleski, 2017; Londo *et al*., 2023). This method uses thermoelectric modules to record voltage changes that are associated with the release of heat that occurs due to the phase change of water into ice (Mills *et al*., 2006). The LTE is associated with the lethal temperature for each sampled primary bud and is correlated with manual assessments of bud death (Wolf and Cook, 1994). Dormant buds were excised from cane tissue leaving an intact bud cushion layer of approximately 1 mm in depth. Buds were placed with the cut-side down on a single ply layer of moistened kimwipe (Kimberly-Clark Corp., USA) to induce ice nucleation during freeze tests (Londo *et al*., 2023). Buds were covered with a square of thermal open-cell foam and sample plates were placed in a programmable freezer (Tenney T2C, Tenney Environmental, New Columbia, PA, USA) connected to a Keithley 2700 or 2701 multimeter datalogger (Keithley Instruments). Voltage and temperature measurements were collected at 15-second intervals using the BudFreezer program (Brock University, Guelph ON, Canada). Freezing runs were conducted as follows: 1°C/minute cooling ramp to 0°C, hold for 1 hour, 4°C/hour cooling ramp to −40°C, hold one hour at −40°C, 4°C/hour warming ramp to room temperature. Low temperature exotherm peaks were manually annotated using BudProcessor/BudLTE software (Brock University, Guelph ON, Canada). In the first year, 2018-2019, 5 replicate buds were assessed for initial LTE measurements. In the second and third years of the study, 10 replicate buds were used for initial LTE measurements. This adjustment was made to increase the confidence in the mean LTE value as representative of field cold hardiness level. Here we report the mean LTE value for any given time point as the LT50.

### Assessment of deacclimation resistance and deacclimation potential

To determine deacclimation resistance of the buds collected during weekly collections, twenty-five single node cuttings were placed in plastic cups with basal cut ends in water. Cups were placed in a Conviron growth chamber (Controlled Environments Inc., North Dakota, USA) and held at constant 20°C without light exposure. Cold hardiness was assessed as described above to measure the loss of cold hardiness under forcing conditions at repeated intervals (e.g. T0-field collected, T1 = T0+6 days, T2 = T0+12 days, ect). Resampling intervals varied throughout the experiments with longer intervals in early winter and shorter intervals in late winter, as deacclimation resistance declined. At each deacclimation time point, five replicate buds for each cultivar were assessed to determine the change in LT50 from the previous time points. Deacclimation rates for each cultivar at each collection time point were analyzed independently using linear regression and taking the slope of the regression as the deacclimation rate (e.g., lm(LTE∼SampleDay). To compare deacclimation rates between cultivars as a function of chilling hour estimates, raw rate values for each year were normalized within cultivars using the mean of the three highest rate values observed across the study. The normalized percentage of total deacclimation is referred to as deacclimation potential (Ψ_deacc_). (Kovaleski *et al*., 2018; Kovaleski, 2022; North *et al*., 2022). The nls() function of the drc package (Ritz *et al*., 2015) was used with default parameters (fct=LL.2(), type=“continuous”) to model Ψ_deacc_ between cultivars, years, and in comparison with the various chill model estimates as described in North (2022).

### Assessment of budbreak phenology

In tandem with field LTE assessments, an additional 10 single node cuttings were placed in plastic cups with basal ends trimmed and kept in water. Cups were left at 18-20°C temperatures and without specific light exposure (laboratory lighting). Budbreak phenology was manually assessed on alternating days during the week. Days to bud break (DTB) was recorded as the number of days between field collection and first evidence of green tissue, EL stage 3 (Coombe, 1995). When a cutting was marked as EL 3, they were removed from the experiment to prevent double counting. Budbreak data was manually curated to remove sample collection points where less than 5 of the 10 replicates achieved budbreak in under 150 days. For the remaining collections, budbreak was phenotyped as the average (meanDTB) time to budbreak.

## Results

### Characterization of the three winters

Temperatures during the three winter seasons of this study were typical for the location. Seasonal minimum temperatures occurred earlier, and was colder, in year 1 (−20.9°C) compared with years 2 (−17.7°C) and 3 (−17°C). The three years of study did not substantially differ in the pattern of chill accumulation modeled, resulting in high correlation between chill accumulation rates for each of the chill models (**Figure S1, Table S2**). Therefore, we restricted our analysis to examining chilling accumulation relative to the Dynamic Portions model. The maximum estimated chilling received at the end of sample collection in each year were broadly similar: Year 1 was terminated on 3 April with 103 portions, Year 2 on 11 March with 91 portions, and Year 3 on 31 March with 98 portions.

### Cold hardiness

Cold hardiness data was collected from all cultivars and all three years with five buds sampled at each of 31 weekly collections in 2018-2019 (n=4,805), and 10 buds sampled at each of 28 or 27 weekly collections in 2019-2020 (n=8,680) and 2020-2021 (n=8,370) respectively. In total, 21,855 buds were sampled for determining field cold hardiness across the three years. Observed DTA peaks for field collected samples represented 84.8% (n=4,073), 72.6% (n=6,304), and 82.9% (n=6,936) of expected peaks in each year respectively. Outliers were removed from LTE data for initial field collections (weekly sampling) and deacclimation data (repeated sampling, see below) based on studentized residuals >3 on linear models for cold hardiness in respect to time. Deacclimation timepoints were removed where not enough LTE peaks were resolved within a collection to plot rate regressions (minimum three time points). In total, 59,001 LTE observations were recorded and of these, 4% of observations (n=2,323) were removed from the study.

Cold hardiness phenotypes (LT50s) followed a similar seasonal pattern with enhancement (acclimation) of cold hardiness in fall and early winter, maintenance during winter, and deacclimation in late winter/early spring (**Figure 1**). While the exact shape of the cold hardiness curve differed by year and cultivar, the average maximum cold hardiness was broadly comparable between years, within each cultivar. As a group, *vinifera* cultivars were less cold hardy than *hybrid* cultivars. Merlot and Syrah were observed to be the least cold hardy with an average maximum LT50s of −21.1°C and −21.6°C respectively. In contrast, Marechal Foch and Concord were observed to be the most cold hardy with LT50 values of −27.6°C and −27.2°C, respectively. Cold hardiness curves and maximum cold hardiness values for all cultivars and all years can be found in **Figure S2** and **Table S3**.

**Figure 1:**
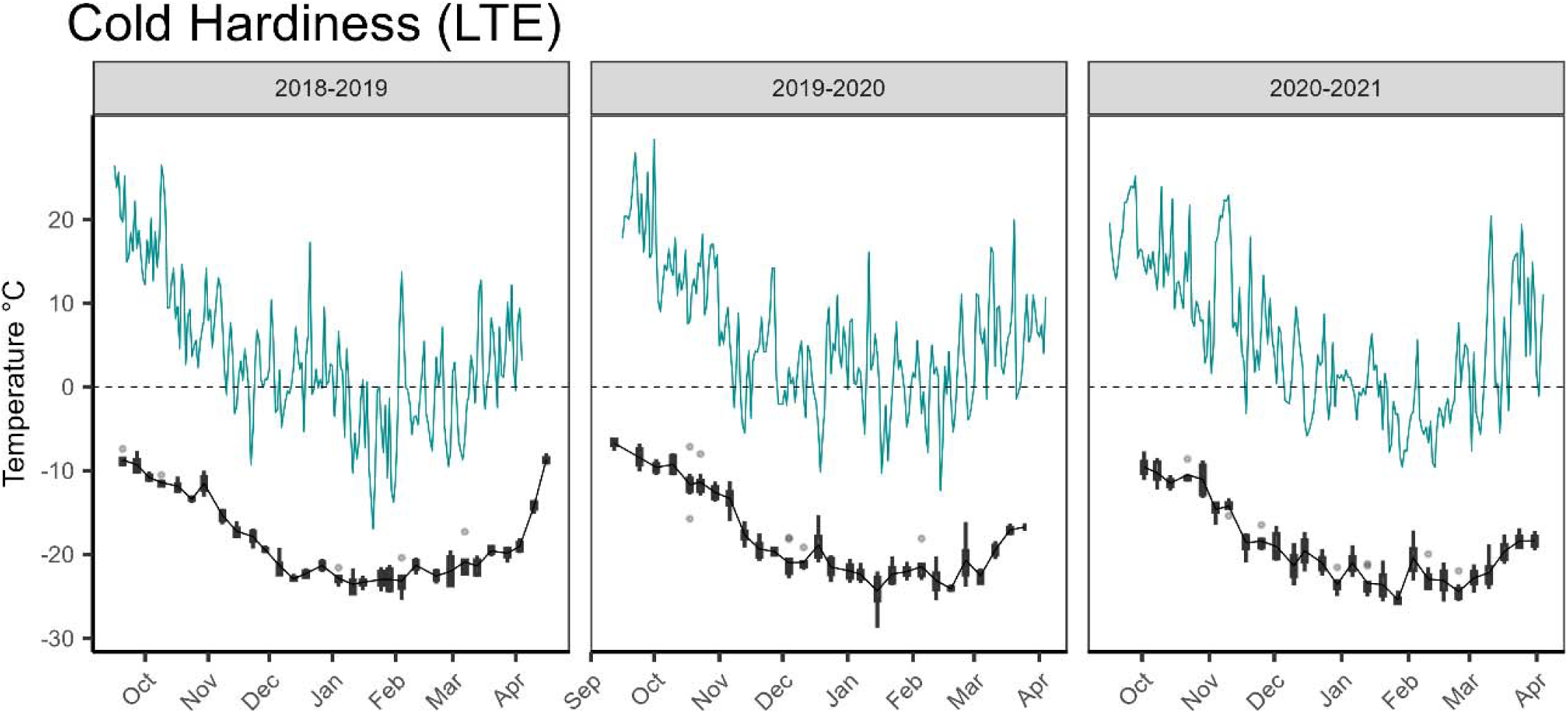
Cold hardiness data for *V. vinifera* ‘Riesling’ across three different winter seasons as assessed with differential thermal analysis. Blue line indicates minimum temperatures in the three years of this study. Black lines and boxplots indicate weekly LT50 values computed from low temperature exotherm data from dormant buds

### Deacclimation rates

Deacclimation rates were determined from the linear relationship of cold hardiness lost with respect to time under forcing for each cultivar at each collection timepoint across the three years of the study (**Table S3**). Deacclimation experiments were disrupted during year 1 due to the 2018-2019 US Federal Government furlough that occurred on 22 December 2018 and extended until 25 January 2019. Experiments initiated just prior to the furlough and through the remainder of the 2018-2019 season had to be moved from the 20°C growth chamber into a similar storage space where temperatures were slightly cooler and averaged 18°C. The lower forcing temperature resulted in lower deacclimation rates in the 2018-2019 dataset when compared with 2019-2020 and 2020-2021, following a previously shown effect of forcing temperature on deacclimation rates (Kovaleski *et al*., 2018). To compensate, deacclimation rates for collections occurring after the disruption in 2018-2019 were scaled by dividing the measured rates by the ratio of the maximum rate in year 1 compared with the averaged maximum rates of years 2 and 3 (**Table S4**). The maximum rates for each year were determined by taking the average of the three highest observed rates in each year and can be found in **Table S5**. The fastest deacclimating genotypes included *hybrid* cultivars Marechal Foch (2.87°C/day), Niagara (2.46°C/day) and Concord (2.29°C/day), and *vinifera* cultivar Gewurztraminer (2.21°C/day). The slowest deacclimating genotypes included *hybrid* cultivars Chambourcin (1.03°C/day) and Vignoles (1.14°C/day) as well as *vinifera* cultivars Tocai Fruliano (1.28°C/day), Syrah (1.37°C/day), and Sauvignon blanc (1.39°C/day). Deacclimation rates changed throughout winter as a function of chilling hour exposure in a log-logistic response (**Figure 2**) and curves for all cultivars can be found in **Figure S2**. For each cultivar, deacclimation potential (Ψ_deacc_) was calculated by normalizing the observed deacclimation rates for each weekly collection in each year relative to the computed max rate and fit to a log-logistic model where the inflection point represents a Ψ_deacc_of 50%. Despite wide variation in maximum deacclimation rates among cultivars, very little variation in Ψ_deacc_was observed (**Figure 2**), with an experiment-wide average Ψ_deacc_ inflection (50% Ψ_deacc_) of 51 Dynamic portions (**Table S6)**.

**Figure 2:**
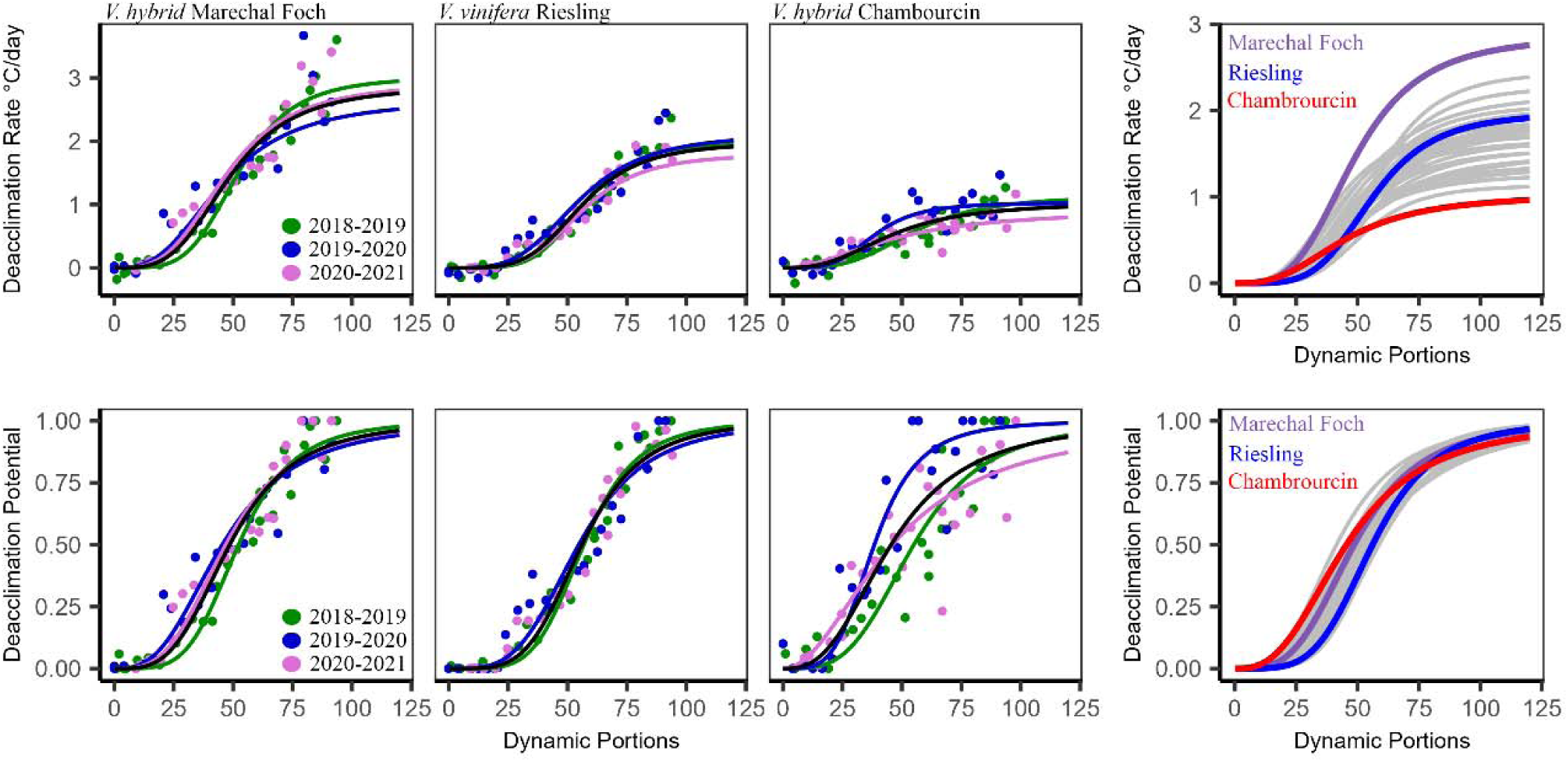
Deacclimation rates and deacclimation potential of three grapevine cultivars, ‘Marechal Foch’, ‘Riesling’, and ‘Chambourcin’. Top panels show measured deacclimation rates in colored points for each year of the study. The mean of the three years is shown in black. Curves are a log logistic fit of the data. Deacclimation rates across years as driven by chilling accumulation are conserved within cultivar but differ dramatically between cultivars. Bottom panels show deacclimation potential,Ψ_deacc_, for each cultivar. Right hand panels show selected cultivar responses relative to the whole dataset in gray.

### Budbreak phenology

Budbreak phenology was recorded as the mean days to budbreak (meanDTB) under forcing conditions. At the start of each season, clear indications of endodormancy were observed, typified by long time to budbreak in all cultivars and low overall percentage of break within the sets of 10 replicate buds (**Figure 3**). As chilling accumulated, all cultivars demonstrated increased synchrony and increased complete break of the replicates (**Figure S2**). Estimates of chilling requirements based on meanDTB were determined using a critical threshold of 28 days (Londo and Johnson, 2014). Cultivars that required the least chilling to reach meanDTB at 28 days were Chardonnay, Gewurztraminer, and Marechal Foch (54, 40, and 39 Portions respectively) while cultivars that required the highest chilling included Traminette, Cabernet Sauvignon and Chambourcin (83, 85, and 94 Portions respectively). Chilling requirements for all cultivars can be found in **Table S7**.

**Figure 3:**
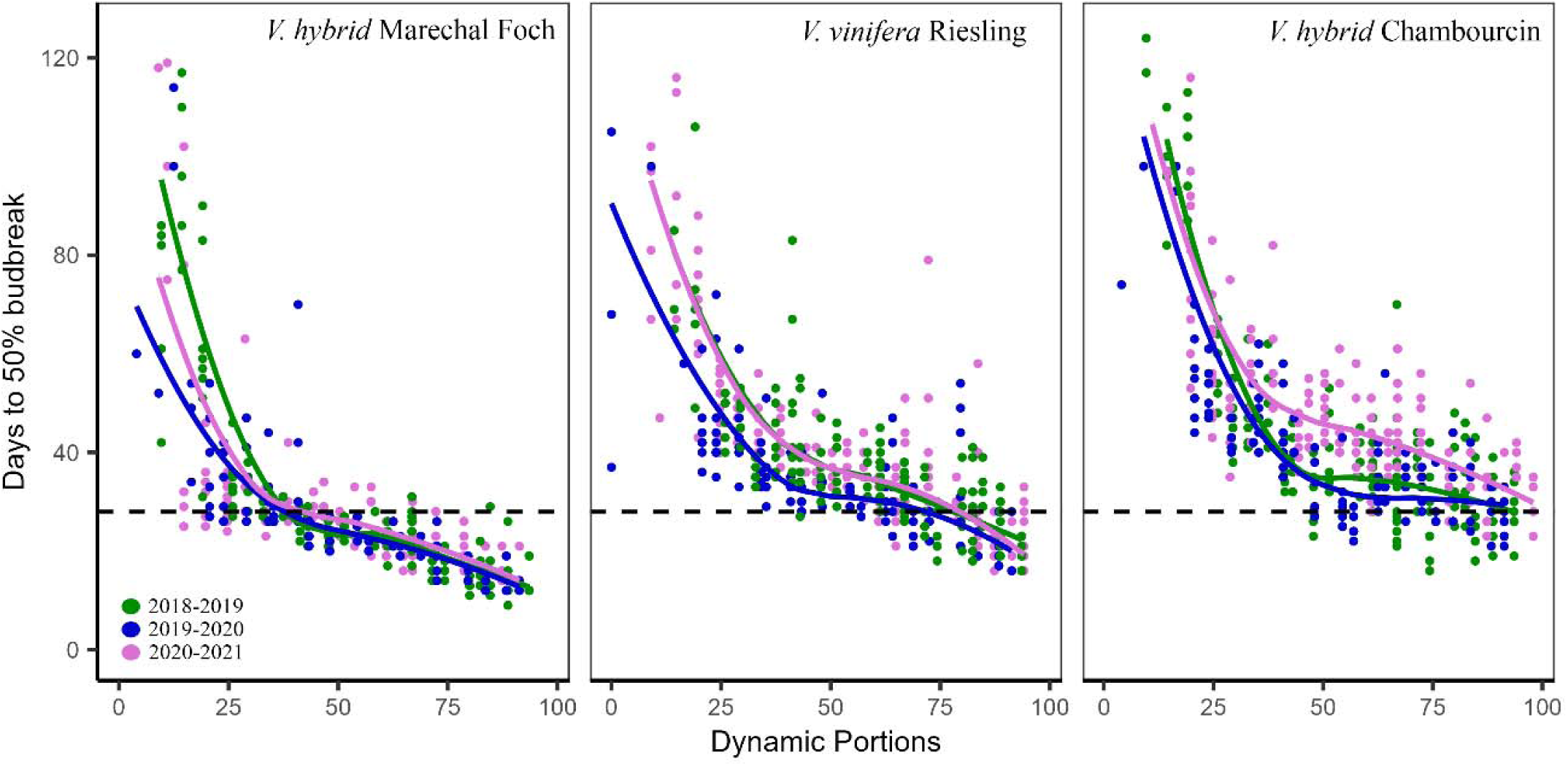
Budbreak response under forcing conditions for cultivars Marechal Foch, Riesling, and Chambourcin. Dashed line indicates 28 days in forcing conditions and the cutoff point for assessing chilling requirement.

### Correlation of winter traits with cultivar type or heritage

Linear models and ANOVA were used to examine the potential for adaptive signatures in winter physiology traits based on cultivar heritage. First, trait values were examined relative to the binary designation of *vinifera* vs. *hybrid* (“Type”). Second, cultivars were separated based on “Heritage”: *Vinifera* cultivars were categorized as “cool” or “warm” cultivars based on growing season responses (Jones *et al*., 2012); *Hybrid* cultivars were partitioned into those with a breeding history that utilized predominantly wild North American grapevine species with northern distributions such as *V. riparia* and *V. labrusca* (“Northern”) or those that utilized wild species with southern distributions such as *V. rupestris* and *V. lincecumii* (“Southern”). Traits evaluated included maximum deacclimation rates observed by the end of each winter, maximum cold hardiness in each year, chilling requirement based on 50% budbreak at 28 days under forcing conditions for each year, and chilling requirement as determined by the inflection point in the Ψ_deacc_ curves for each year (**Figure 4**). Differences based on “Type” were observed for maximum deacclimation rate (p=0.08) with *hybrid* cultivars having slightly faster average deacclimation rates, and for cold hardiness (p<0.001), where hybrids as a group had a significantly deeper maximum cold hardiness. No significant differences were observed between *vinifera* and *hybrid* for chilling requirement based on forcing assays (p=0.57) nor for chilling requirement based on deacclimation potential (p=0.82) (**Table S8**).

**Figure 4:**
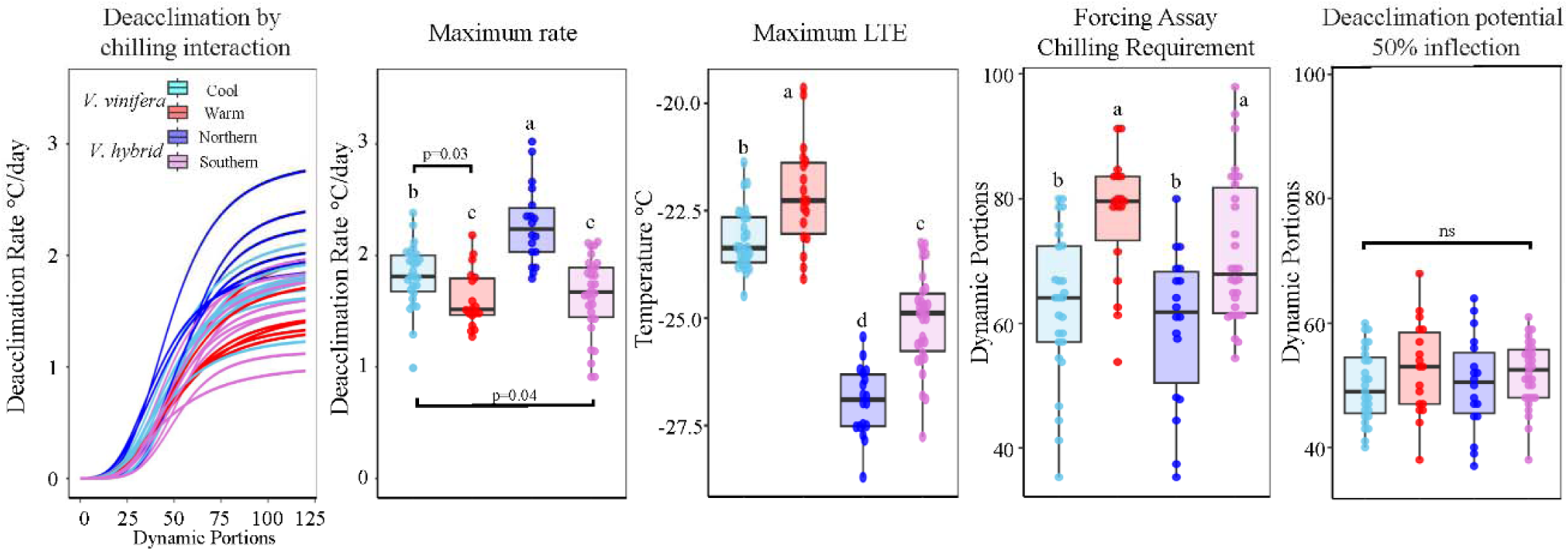
Relationship of winter physiology traits with grapevine cultivars type and heritage. First panel shows the relationship of chilling accumulation as measured by the Dynamic Portions model with deacclimation rate under forcing conditions. Curves represent the average of all three years for cultivar response. Panel two-five shows significant differences between heritage classes of grapevine for maximum rate, maximum LTE, and chilling requirement as determined by forcing assays. Deacclimation potential does not significantly differ.

When cultivars were contrasted based on “Heritage”, significant differences were observed for maximum deacclimation (p<0.001), maximum cold hardiness (p<0.001), and chilling requirement based on forcing assays (p<0.001), but not for chilling requirement based on deacclimation potential (p=0.42) (**Table S8**). Northern *hybrids* had the highest deacclimation rates, followed by cool climate *vinifera*. Warm climate *vinifera* and Southern *hybrids* had the lowest deacclimation rates. For maximum cold hardiness, all groups were significantly different from each other, where warm climate and Southern genotypes were less cold hardy than their within Type counterparts. For chilling requirement as determined by forcing assays, warm climate *vinifera* and Southern *hybrid* classes needed significantly greater chilling to reach budbreak compared with cool climate *vinifera* and Northern *hybrids* (**Figure 4**).

### Linking cold hardiness, deacclimation, and budbreak

To test the hypothesis that cold hardiness (LT50) and deacclimation rate are mechanistically linked to budbreak phenology, we examined the potential to predict budbreak in assays using deacclimation dynamics. To do this, we used the equation: Predicted budbreak=|LT50 - CH_BB_|/Deacclimation rate, where CH_BB_ is the cold hardiness at budbreak. Here, rather than direct estimates of cold hardiness for CH_BB_ based on deacclimation assays (as in Kovaleski 2022), we estimated CH_BB_ based on 1°C iterations between –5°C and 20°C for each cultivar, evaluated based on RMSE. Positive values of cold hardiness for CH_BB_ are possible as they just denote a thermal time measurement once cold hardiness crosses tissues threshold (∼-2 °C). Predicted budbreak was regressed against mean observed time to budbreak applying three levels of filtering to the data: Ψ_deacc_>10%, Ψ_deacc_>50% and Ψ_deacc_>75%. RMSE values were highest when data representing low levels of deacclimation potential were included (10% filter), and lowest with higher Ψ_deacc_ (75% filter), denoting our best ability to predict budbreak occurs at higher chill accumulation (**Figure 5**, **Table S9**). Cultivar specific values of CH_BB_ with lowest RMSE were generally positive (8°C to 20°C) and the experiment average CH_BB_ was 17°C. Plots of predicted versus observed days to budbreak for all cultivars at all CH_BB_ iterations can be found in **Figure S3**.

**Figure 5:**
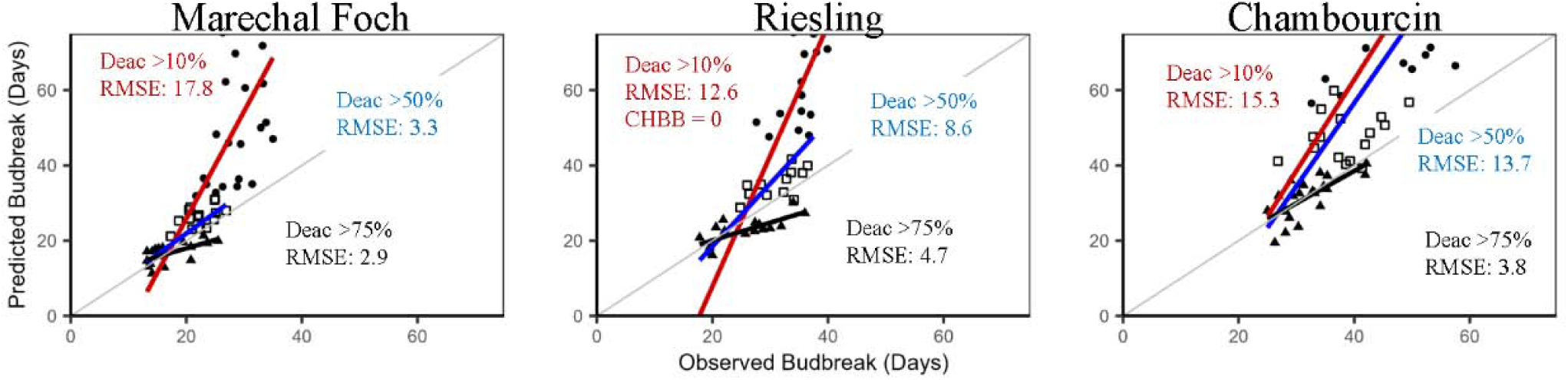
Relationship between predicted days to budbreak, determined by integrating LTE and deacclimation rate data for each collection time point, vs observed budbreak in forcing conditions. Point shapes indicate chilling accumulation in Dynamic portions for weekly collections from across the three years of the study. Points and regression shown for the CH_BB_ with the lowest RMSE for each cultivar: Marechal Foch and Riesling CH_BB_ = 20°C, Chambourcin CH_BB_ = 8°C. Red line:black circle indicates the relationship for all collections where Ψ_deacc_ > 10%, blue line: open square for Ψ_deacc_>50% (inflection point), and black line: closed triangle Ψ_deacc_>75%.

## Discussion

Winters are warming faster than any other season in the continental United States, but the implications of disrupted winter physiology extends into the growing season (Burakowski *et al*., 2022). While rising minimum temperatures may allow growers to utilize more cold sensitive genotypes, warmer and less stable winters pose a significant threat to longer-term sustainability of perennial crop sysetms due to false springs and frost risk. The goals of this study were to comprehensively evaluate winter physiology traits across a broad selection of grapevine cultivars to understand if chilling requirements differ between *vinifera* and *hybrid* genotypes, if *hybrid* cultivars are at greater risk from warming winters, and how annual variation in cold hardiness interacts with chilling accumulation and impacts budbreak phenology. Our results demonstrate that as a group, *hybrid* cultivars are more cold hardy than the *vinifera* cultivars – a testament to the successful introgression of this trait into cultivated germplasm by breeding programs. Results also illuminate a potentially maladaptive trait in some cool climate *vinifera* and *hybrid* cultivars: their rapid deacclimation response. Finally, this study validated a link between cold hardiness and deacclimation rates as it relates to budbreak phenology, revealing the potential to predict budbreak with cultivar specific thresholds.

Grapevine cold hardiness has long been a topic of study (as reviewed by (De Rosa *et al*., 2021). *Hybrid* cultivars typically have greater cold hardiness than *vinifera* cultivars and the magnitude of differences depends on the severity of winter temperatures, though studies typically compare only a few varieties at any one location (Ferguson *et al*., 2011, 2014; Londo and Kovaleski, 2017; North *et al*., 2021). In this study, the winter conditions were roughly similar across all three years, and we saw clear differentiation in cold hardiness capacity among the sampled cultivars. *V. vinifera* cultivars as a group were significantly less cold hardy than *hybrid* cultivars, though the range of variation overlapped with the least hardy *hybrid* Cayuga White (−23.4°C) overlapping with the hardiest *vinifera* Riesling (−23.9°C), Lemberger (−23.7°C), Pinot noir (−23.5°C) and Chardonnay (−23.4°C). Cold hardiness variation between the most and least cold hardy cultivars differed by 6.8°C, 5.3°C, and 9.0°C in the three years of this study (**Table S3**), demonstrating wide variation in phenotypic expression and the genetic potential in this trait.

Chilling requirement (CR) is typically evaluated using budbreak forcing assays and chilling is considered fulfilled when buds break synchronously or by a threshold time in forcing. Here we used the latter, using a threshold of 28 days to contextualize our dormancy evaluation results. Wide variation in CR among cultivars was observed when using this metric, ranging from a low chill minimum of 39.0 portions for the *hybrid* Marechal Foch to a high chill maximum of 94.3 portions for *hybrid* Chambourcin (a difference between mid-December to mid-March in the years observed (**Figure S1**)). While the distribution of CR in this study was continuous, we classified cultivars into 3 CR categories based on equal breaks in the distribution: Low = 40-60, Medium = 60-80, High = 80-100. Examples of low, medium, and high CR were observed both for *vinifera* and *hybrid* cultivars, indicating that CR differences are not solely due to cultivar type (**Figure 6**). Our classification broadly agrees with other studies.

**Figure 6:**
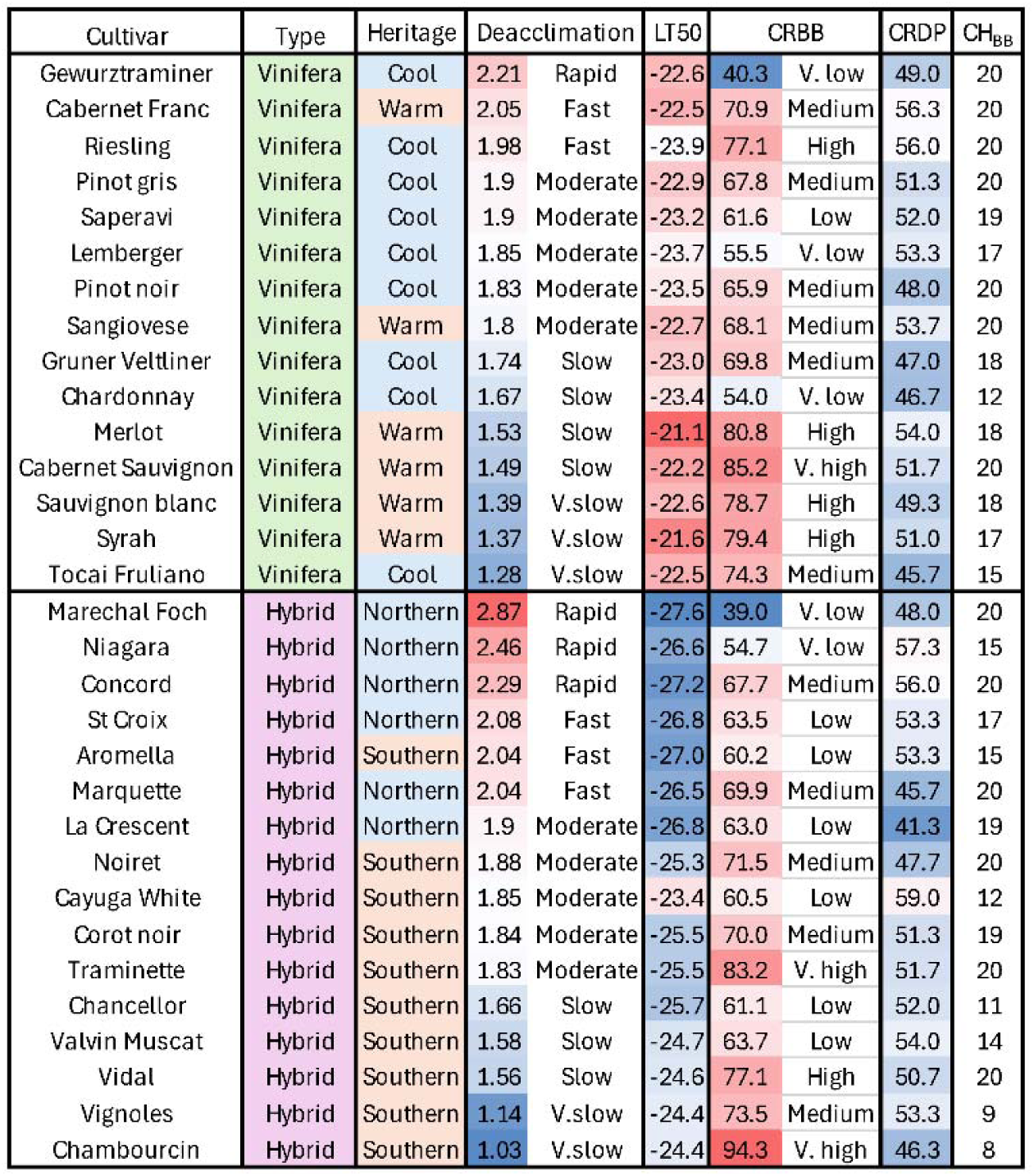
Variation in winter physiology traits measured in this study sorted on cultivar type and deacclimation rate values. Deacclimation rates shown as the average maximum rate in °C/day. Cold hardiness shown as average of maximum cold hardiness as determined by LT50 in °C. Chilling requirement as estimated with forcing assays (CH_BB_) and classification of chill shown as Dynamic portions. Chilling requirement as determined with 50% Ψ_deacc_ (CRDP) shown as Dynamic portions. Temperature parameter for predicting budbreak using cold hardiness and deacclimation rates (CH_BB_).

Comparisons of CR using cumulative chilling hours in Southern Brazil showed Chardonnay, Merlot, and Cabernet Sauvignon required 136, 298, and 392 hours respectively (Anzanello *et al*., 2018), compared to our study which found 54, 80.8, and 85.2 dynamic portions. Similarly, our previous work found CR using the Utah model for Marechal Foch (<250 hours), Concord (1250h), Cabernet Franc (750h), Riesling (750h) and Cabernet Sauvignon (1500h) (Londo and Johnson, 2014), compared to this study which found 39.0, 67.7, 70.9, 77.1, and 85.2 dynamic portions respectively for those cultivars. Despite the variations in chilling accumulation that exists in comparisons across regions (Luedeling and Brown, 2011; Fernandez *et al*., 2020), the rank orders in CR observed are very similar across these studies. Climate heritage significantly affected CR, where paradoxically the genotypes that generally experience longer winters (cool climate *vinifera* and Northern *hybrids*) required less chill exposure to reach budbreak thresholds relative to those from shorter winters (warm climate *vinifera* and southern pedigree *hybrids*). This suggests some maladaptation relative to chilling requirements as currently understood. Colder climate genotypes become ecodormant early in winter, only relying on low temperatures to avoid budbreak. Rising winter temperatures and increased frequency of false springs (Rochette *et al*., 2004; Gu *et al*., 2008; Augspurger, 2013) will thus result in advancements in spring phenology in temperate climates (Chuine *et al*., 2016; Chamberlain and Wolkovich, 2021; Mo *et al*., 2024) and presumably higher frost risk.

Deacclimation assays revealed that early winter field collections displayed very low responsiveness to forcing temperatures, with measured deacclimation rates often negative or not significantly different from zero (**Table S4**). As chilling exposure increased, deacclimation rates gradually shifted from zero to positive values. This transition began slowly, accelerated in midwinter, and peaked by the end of the season (**Figure 2**). Maximum deacclimation rates varied among cultivars (1.03 - 2.87°C/day), with *hybrids* displaying both the highest and lowest rates, while *vinifera* cultivars fell in between (**Figure 4**, **Figure 6**). Notably, our findings reveal a connection between the pedigree of *hybrid* grapevines and the provenance of *vinifera* in relation to deacclimation response. Specifically, *hybrids* selected for deep midwinter cold hardiness exhibited the highest maximum deacclimation rates. These cultivars such as Marchel Foch, Marquette, Niagara, and Concord include northern wild grape species like *V. riparia* and *V. labrusca* in their breeding backgrounds. Similarly, “cool climate” *vinifera* cultivars such as Gewurztraminer, Riesling, and Pinot noir also show faster maximum deacclimation rates. These trends further support a potential adaptive relationship between cool climate suitability and rapid loss of cold hardiness in late winter. We hypothesize that rapid deacclimation may have been beneficial toward grapevine fitness for genotypes with short growing seasons through facilitating loss of deep midwinter cold hardiness in the spring and advancing bubreak. Consequently, it is likely that as climate change warms winters, reducing the need for deep midwinter hardiness and simultaneously advancing late winter deacclimation, frost risk may increase for Northern pedigree *hybrid* varieties and cool climate *vinifera*.

Southern *hybrids* and Warm climate *vinifera* have less cold hardiness and lower deacclimation rates compared to their “heritage” counterparts. *Hybrid* cultivars with slow and very slow deacclimation responses prominently feature *V. rupestris* and *V. lincecumii* in their pedigrees as key wild species. These wild species are native to more southern regions of the United States and while they have sufficient cold hardiness to survive winter conditions in New York (Londo and Kovaleski, 2017), appear to have reduced deacclimation response (Kovaleski and Londo, 2019). Similarly, among the *vinifera* cultivars in our dataset with slower deacclimation rates, most are considered warm or hot climate cultivars (e.g. Merlot, Syrah, Cabernet Sauvignon). Slow deacclimation responses in warm climates would presumably protect grapevines from midwinter warming and early spring frost events through retaining cold hardiness and delaying field budbreak. The physiological and genetic mechanisms that convey rapid or slow deacclimation responses are unknown, but they may also impact the timing of subsequent growing season phenotypes (Sandor *et al*., 2021) as has been shown in Mediterranean species (Segrestin *et al*., 2018). Assessments of growing season phenology in *vinifera* demonstrate phenophases are strongly correlated (Jones and Davis, 2000), with potential to shift in response to climate (Cameron *et al*., 2022). Similarly, wild and hybrid grapevines which have early budbreak also tend to flower, fruit and ripen early, while those with late budbreak have delayed responses (Gutierrez *et al*., 2021). Our survey of cultivars was broad, but is restricted to regionally important cultivars. Future studies evaluating a larger complement of cultivated grapevines from a larger distribution of regional climates would enable a stronger test of the adaptive hypothesis of deacclimation rate proposed here.

The data collected in this study enable us to extrapolate new meaning for chilling requirements within *Vitis*. Our results confirm that chilling requirements based on forcing assays, but also maximum deacclimation rates, are cultivar specific and may indicate climate adaptation in grapevine for winter physiology. Integration of these phenotypes enable two critical advances in our understanding of winter physiology in grapevine, 1) our interpretation of chilling requirement in grapevine requires reevaluation in light of conserved deacclimation potential and 2) understanding of cold hardiness dynamics can contribute to our ability to predict budbreak. Deacclimation rates for all cultivars in this study changed in a logistic pattern, with near zero rates in early winter, a rapid increase in rates in midwinter, and a flattening of maximum rates by the end of winter. While absolute deacclimation rates varied widely among cultivars, normalizing these rates as Ψ_deacc_revealed a consistent, near-universal transition throughout winter, supporting an updated conceptual framework for chilling requirement (**Figure 2, Figure S2**). These results support prior observations that the pattern and timing of dormancy progression in woody perennials is a conserved trait, and differences in chilling requirement are evidence of variation in deacclimation rate (Kovaleski *et al*., 2018; Kovaleski, 2022; North *et al*., 2022). Therefore, while chilling requirements can be used to evaluate local adaptation of cultivars, they do not describe a specific transition in dormancy state. Our current understanding of chilling requirements would indicate no physiological changes in dormancy after the chilling requirement is met. Instead, we observe that deacclimation rates continue to increase after those thresholds are met. The deacclimation rate differences of cultivars then manifest in perceived differences in chilling requirement. High chill requirement (slow deacclimation) cultivars such as Cabernet Sauvignon or Chambourcin do not need longer winters to transition dormancy states compared to low chill cultivars: they are always slow and will always break after a low chill (fast deacclimation) cultivar like Gewurztraminer or Concord. This may seem a semantic difference, but it may be the nuance required to fully understand dormancy physiology through comparisons of high and low chill cultivars of grape, and other fruit crop species. For example, contrasting the transcriptome or metabolome of a high chill and low chill cultivar at timepoints in early winter (endodormancy) or late winter (ecodormancy) across cultivars with the goal of understanding the mechanisms of chilling requirement would not contrast differences in dormancy status. Instead this contrast would be comparing the cultivar level variation in deacclimation rates of these two cultivars at the same physiological timepoints of ∼25% and ∼75% of deacclimation potential. Candidate gene or candidate metabolite selection in these contrasts would potentially identify growth response genes, rather than true chilling hour related genes.

Besides implications on our understanding of dormancy, another goal of this study was to evaluate the range of variation in cold hardiness and deacclimation response in grapevine, and how these traits can be used to predict budbreak. Phenology modeling has been successful for many crop species and for many growth stages (Parker *et al*., 2011, 2013; García de Cortázar-Atauri *et al*., 2017), yet modeling budbreak in perennial species remains a challenge, particularly due to interactions between and timing of chill and heat accumulation (Chuine *et al*., 2016). While there are many different models for grapevine budburst (García de Cortázar-Atauri *et al*., 2009; Nendel, 2010; Wang *et al*., 2020; Piña-Rey *et al*., 2021; Pellegrino *et al*., 2022), these models can underperform when compared across years or across regions. One of the likely explanations for this challenge is that budbreak prediction has long relied on the concept of growing degree days (GDD) (García de Cortázar-Atauri *et al*., 2009), the heat accumulation that occurs after a somewhat arbitrary start point, typically after January 1st in the Northern Hemisphere. We propose that one of the reasons for the underperformance of GDD estimates for budbreak is these estimates do not account for variation in cold hardiness and deacclimation rate. Recently a mechanistic link between cold hardiness and deacclimation rate was used to predict field budbreak phenology among diverse woody species (Kovaleski, 2022). Applying this framework to grapevine budbreak in controlled environment demonstrated a clear association of these traits with budbreak, but prediction accuracy was initially poor when using data from the entire season (**Figure S3**). Regressions that included field collections with low Ψ_deacc_ (or low chill accumulation) were skewed towards a delayed prediction of budbreak. This suggests that measured deacclimation rates for early winter were underestimated or that budbreak phenology as measured by meanDTB does not describe phenology well (requiring some incorporation of budbreak distribution such as in (Camargo Alvarez *et al*., 2018)). Given the differences in bud internal bud morphology between wild and *vinifera* grapes (Kovaleski *et al*., 2019), it is likely that variation in bud swell progression and eventual separation of the bud scales to reveal green bud tissue (EL stage 3) impact the consistency of budbreak quantification across species and interspecific hybrids. Restricting regressions to collection timepoints that correspond to the cultivar wide inflection point in Ψ_deacc_ (>50%) or greater (>75%) greatly reduced error in the prediction (**Table S9**). In this study the midpoint in Ψ_deacc_ occurs at 51 Dynamic portions and corresponds to the end of December in each of the three study years (12/26/2018, 12/29/2019, 12/23/2020). As the Ψ_deacc_response was conserved among cultivars, this result demonstrates that there is some utility in the concept of using GDD after January 1 as a component of budbreak prediction. Our results show that by incorporating chilling accumulation and cold hardiness status from Jan 1 onward, either from direct field measures with DTA or through using validated cold hardiness models (Ferguson *et al*., 2014; North *et al*., 2021; Kovaleski *et al*., 2023; Wang *et al*., 2023), could be sufficient to predict budbreak. An important limitation of our results is that we measured deacclimation responses and predict budbreak only at 20°C, which is much warmer than typical spring temperatures. Previous research has demonstrated that the impact of temperature on deacclimation rate in grapevines is not linear, particularly demonstrating exponential increases at low temperatures (Kovaleski *et al*., 2018); North *et al*., 2022). It is possible that predicting budbreak for specific grapevine cultivars may require evaluation of deacclimation response at lower temperatures, and validations of field phenology prediction may need to include diurnal temperature fluctuations rather than daily means.

## Conclusion

This study describes a comprehensive examination of key winter physiology traits in multiple *hybrid* and *vinifera* grapevines. We demonstrate support for a hypothesis that while genetic differences explain variation in maximum cold hardiness between *hybrid* and *vinifera* cultivars, deacclimation rates appear to be adaptive within type, depending on climate of origin. Correlation between deacclimation and budbreak phenology with cultivar heritage, demonstrates a predisposition of northern *hybrid* and some cool climate *vinifera* to greater frost risk as climate warms. However, southern wild and *hybrid* grapevines may offer new important germplasm for developing climate resilient grapevines through the breeding of slow deacclimating, frost avoiding cultivars. We also demonstrated through examining deacclimation potential that the progression of all grapevine cultivars through dormancy is conserved, though heat efficiency is cultivar specific. This important nuance changes our framework for the concept of chilling requirement and cultivar suitability that should be considered when conducting physiological and -omics based analysis of the dormancy and chilling accumulation mechanism. Finally, we show that budbreak phenology is in part mechanistically determined by the interaction of dormant bud cold hardiness and deacclimation rate response. Combining these traits, we identified that the critical budbreak parameter CH_BB_ varies across genotypes and can be used to predict budbreak in grapevine. Validation in the effectiveness of these predictions for field budbreak relative to other prediction models remain, but it is clear that cold hardiness and deacclimation rate traits play a critical role in understanding how climate change will impact phenology.

## Acknowledgement

We thank Hanna Martens, Felex Pike, and Kathleen Deys for help collecting and processing grape bud tissue for cold hardiness measurements, sampling for LTE, and budbreak phenotyping. We would also like to thank the NE1020 experimental vineyard, Ravines, Anthony Road, Swedish Hill, Ventosa, Three Brother’s, Wagner, PreJean, Wiemer, Standing Stone, and Seneca Shore Vineyards for access to *V. vinifera* and *V. hybrid* vines. This work was partially supported by the Office of the Vice Chancellor for Research and Graduate Education at the University of Wisconsin–Madison with funding from the Wisconsin Alumni Research Foundation, through the USDA-ARS appropriated project 1910–21220–006–00D, the USDA National Institute for Food and Agriculture (award: 2023-68008-39274) and the New York Wine and Grape Foundation.

## Data Availability

Data for cold hardiness, budbreak, and deacclimation available upon request or via open access at Figshare:

https://doi.org/10.6084/m9.figshare.27084265.v1

https://doi.org/10.6084/m9.figshare.27084268.v1

https://doi.org/10.6084/m9.figshare.27084271.v1

---

Figure S1: Comparison of temperature conditions over the three years of the study. Different models for measuring chill accumulation shown as overlapping temperature ranges. Accumulation of chill across the three winters shown in panel subsets below for each chilling model. Dynamic Portions were used for analysis in this study.

Figure S2: Summary of winter physiology trait responses for all cultivars tested in this study. Top panel shows cold hardiness values across the three winters, represented as LT50 values. Center panels show measured deacclimation rate data with log-logistic curve fit for each year and the average of all years combined, normalization of data as deacclimation potential, and budbreak forcing assays. Bottom panels show deacclimation rate and deacclimation potential for each cultivar relative to the responses measured for all cultivars across the study.

Figure S3: Summary of permuted CH_BB_ values for each cultivar tested in this study. Regression fits for various filtered datasets shown in red (Ψ_deacc_>10%), blue (Ψ_deacc_>50%), and black (Ψ_deacc_>75%). RMSE values for each permuted CH_BB_ can be found in Supplemental Table 9.

Table S1: Grapevine cultivars collected and locations of commercial or experimental vineyards

Table S2: Correlation between chill models

Table S3: Average maximum cold hardiness as determined by the average of the three lowest LT50 values in each year.

Table S4: Deacclimation rates and conversion factor used to adjust for furlough disruption.

Table S5: Maximum deacclimation rate data observed in this study computed as loss of cold hardiness in °C/day at 20 °C.

Table S6: Log-logistic output for cultivar specific deacclimation potential. E parameter specifies the inflection of the curve at 50% of deacclimation potential.

Table S7: Chilling requirement for all cultivars as determined by forcing assays.

Table S8: Anova results of winter physiology traits contrasting type and heritage

Table S9: Cultivar specific parameters for predicting budbreak using LTE and deacclimation response. CH_BB_ indicates the prediction value at the end of the deacclimation vector.

## References

Anderson JV, Horvath DP, Chao WS, Foley ME. 2010. Bud Dormancy in Perennial Plants: A Mechanism for Survival. Topics in Current Genetics. Dormancy and Resistance in Harsh Environments. Springer, Berlin, Heidelberg, 69–90.

Andrews PK, Sandidge CR, Toyama TK. 1984. Deep Supercooling of Dormant and Deacclimating Vitis Buds. American journal of enology and viticulture 35, 175–177.

Anzanello R, Fialho FB, Santos HP dos. 2018. Chilling requirements and dormancy evolution in grapevine buds. Ciência e Agrotecnologia 42, 364–371.

Anzanello R, Fogaça CM, Sartori GBD. 2021. Induction and overcoming of dormancy of grapevine buds in response to thermal variations in the winter period. Ciencia rural 51, e20200887.

Atkinson CJ, Brennan RM, Jones HG. 2013. Declining chilling and its impact on temperate perennial crops. Environmental and experimental botany 91, 48–62.

Augspurger CK. 2013. Reconstructing patterns of temperature, phenology, and frost damage over 124 years: spring damage risk is increasing. Ecology 94, 41–50.

Bigler C, Bugmann H. 2018. Climate-induced shifts in leaf unfolding and frost risk of European trees and shrubs. Scientific reports 8, 9865.

Burakowski EA, Contosta AR, Grogan D, Nelson SJ, Garlick S, Casson N. 2022. Future of winter in northeastern north America: Climate indicators portray warming and snow loss that will impact ecosystems and communities. Northeastern naturalist 28, 180–207.

Camargo Alvarez H, Salazar-Gutiérrez M, Zapata D, Keller M, Hoogenboom G. 2018. Time-to-event analysis to evaluate dormancy status of single-bud cuttings: an example for grapevines. Plant methods 14, 94.

Cameron W, Petrie PR, Barlow EWR. 2022. The effect of temperature on grapevine phenological intervals: Sensitivity of budburst to flowering. Agricultural and forest meteorology 315, 108841.

Campoy JA, Ruiz D, Egea J. 2011. Dormancy in temperate fruit trees in a global warming context: A review. Scientia horticulturae 130, 357–372.

Chamberlain CJ, Wolkovich EM. 2021. Late spring freezes coupled with warming winters alter temperate tree phenology and growth. The new phytologist 231, 987–995.

chillR Package: Statistical Methods for Phenology Analysis in Temperate Fruit Trees. 2022. https://www.rdocumentation.org/packages/chillR/versions/0.75. Accessed September 2024.

Chuine I, Bonhomme M, Legave J-M, García de Cortázar-Atauri I, Charrier G, Lacointe A, Améglio T. 2016. Can phenological models predict tree phenology accurately in the future? The unrevealed hurdle of endodormancy break. Global change biology 22, 3444–3460.

Coombe BG. 1995. Growth Stages of the Grapevine: Adoption of a system for identifying grapevine growth stages. Australian journal of grape and wine research 1, 104–110.

Delgado A, Dapena E, Fernandez E, Luedeling E. 2021. Climatic requirements during dormancy in apple trees from northwestern Spain – Global warming may threaten the cultivation of high-chill cultivars. European journal of agronomy: the journal of the European Society for Agronomy 130, 126374.

Dennis FG. 2003. Problems in standardizing methods for evaluating the chilling requirements for the breaking of dormancy in buds of woody plants. HortScience: a publication of the American Society for Horticultural Science 38, 347–350.

De Rosa V, Vizzotto G, Falchi R. 2021. Cold hardiness dynamics and spring phenology: Climate-driven changes and new molecular insights into grapevine adaptive potential. Frontiers in plant science 12, 644528.

Dokoozlian NK. 1999. Chilling temperature and duration interact on the budbreak of ‘Perlette’ grapevine cuttings. HortScience: a publication of the American Society for Horticultural Science 34, 1–3.

Drogoudi P, Cantín CM, Brandi F, et al. 2023. Impact of chill and heat exposures under diverse climatic conditions on peach and nectarine flowering phenology. Plants 12, 584.

Erez A, Fishman S, Linsley-Noakes GC, Allan P. 1990. The dynamic model for rest completion in peach buds. Acta horticulturae, 165–174.

FAOSTAT. https://www.fao.org/faostat/en/#data/QCL/visualize. Accessed September 2024.

Fennell A. 2004. Freezing Tolerance and Injury in Grapevines. Journal of Crop Improvement 10, 201–235.

Ferguson JC, Moyer MM, Mills LJ, Hoogenboom G, Keller M. 2014. Modeling Dormant Bud Cold Hardiness and Budbreak in Twenty-Three Vitis Genotypes Reveals Variation by Region of Origin. American journal of enology and viticulture 65, 59–71.

Ferguson JC, Tarara JM, Mills LJ, Grove GG, Keller M. 2011. Dynamic thermal time model of cold hardiness for dormant grapevine buds. Annals of botany 107, 389–396.

Fernandez E, Mojahid H, Fadón E, et al. 2023. Climate change impacts on winter chill in Mediterranean temperate fruit orchards. Regional environmental change 23, 1–18.

Fernandez E, Whitney C, Luedeling E. 2020. The importance of chill model selection — a multi-site analysis. European journal of agronomy: the journal of the European Society for Agronomy 119, 126103.

Fraga H, Santos JA. 2021. Assessment of Climate Change Impacts on Chilling and Forcing for the Main Fresh Fruit Regions in Portugal. Frontiers in plant science 12, 689121.

García de Cortázar-Atauri I, Brisson N, Gaudillere JP. 2009. Performance of several models for predicting budburst date of grapevine (Vitis vinifera L.). International journal of biometeorology 53, 317–326.

García de Cortázar-Atauri I, Duchêne E, Destrac-Irvine A, Barbeau G, De Rességuier L, Lacombe T, Parker AK, Saurin N, Van Leeuwen C. 2017. Grapevine phenology in France: from past observations to future evolutions in the context of climate change. OENO One 51, 115–126.

Garris A, Clark L, Owens C, McKay S, Luby J, Mathiason K, Fennell A. 2009. Mapping of photoperiod-induced growth cessation in the wild grape Vitis riparia. Journal of the American Society for Horticultural Science. American Society for Horticultural Science 134, 261–272.

Govaerts R. 2024. Plants of the World Online. Facilitated by the Royal Botanic Gardens, Kew. https://powo.science.kew.org/. Accessed September 2024.

Grassi F, De Lorenzis G. 2021. Back to the origins: Background and perspectives of grapevine domestication. International journal of molecular sciences 22, 4518.

Gu L, Hanson PJ, Kaiser DP, Yang B, Nemani R, Pallardy SG, Meyers T. 2008. The 2007 Eastern US Spring Freeze: Increased Cold Damage in a Warming World? Bioscience 58, 253–262.

Gutierrez B, Schwaninger H, Meakem V, Londo J, Zhong G-Y. 2021. Phenological diversity in wild and hybrid grapes (Vitis) from the USDA-ARS cold-hardy grape collection. Scientific reports 11, 24292.

Hannah L, Roehrdanz PR, Ikegami M, Shepard AV, Shaw MR, Tabor G, Zhi L, Marquet PA, Hijmans RJ. 2013. Climate change, wine, and conservation. Proceedings of the National Academy of Sciences of the United States of America 110, 6907–6912.

Horvath DP, Anderson JV, Chao WS, Foley ME. 2003. Knowing when to grow: signals regulating bud dormancy. Trends in plant science 8, 534–540.

Jones GV, Davis RE. 2000. Climate influences on grapevine phenology, grape composition, and wine production and quality for Bordeaux, France. American journal of enology and viticulture 51, 249–261.

Jones KS, McKersie BD, Paroschy J. 2000. Prevention of ice propagation by permeability barriers in bud axes of Vitis vinifera. Canadian journal of botany. Journal canadien de botanique 78, 3–9.

Jones GV, Reid R, Vilks A. 2012. Climate, grapes, and wine: Structure and suitability in a variable and changing climate. The Geography of Wine. Dordrecht: Springer Netherlands, 109–133.

Kalberer SR, Wisniewski M, Arora R. 2006. Deacclimation and reacclimation of cold-hardy plants: Current understanding and emerging concepts. Plant science: an international journal of experimental plant biology 171, 3–16.

Keller M. 2023. Climate change impacts on vineyards in warm and dry areas: Challenges and opportunities: From the ASEV climate change symposium part 1 - viticulture. American journal of enology and viticulture 74, 0740033.

Kovaleski AP. 2022. Woody species do not differ in dormancy progression: Differences in time to budbreak due to forcing and cold hardiness. Proceedings of the National Academy of Sciences of the United States of America 119, e2112250119.

Kovaleski AP, Londo JP. 2019. Tempo of gene regulation in wild and cultivated Vitis species shows coordination between cold deacclimation and budbreak. Plant science: an international journal of experimental plant biology 287, 110178.

Kovaleski AP, Londo JP, Finkelstein KD. 2019. X-ray phase contrast imaging of Vitis spp. buds shows freezing pattern and correlation between volume and cold hardiness. Scientific reports 9, 14949.

Kovaleski AP, North MG, Martinson TE, Londo JP. 2023. Development of a new cold hardiness prediction model for grapevine using phased integration of acclimation and deacclimation responses. Agricultural and Forest Meteorology 331, 109324.

Kovaleski AP, Reisch BI, Londo JP. 2018. Deacclimation kinetics as a quantitative phenotype for delineating the dormancy transition and thermal efficiency for budbreak in Vitis species. AoB plants 10, ly066.

Lamichhane JR. 2021. Rising risks of late-spring frosts in a changing climate. Nature climate change 11, 554–555.

Lang GA, Early JD, Martin GC, Darnell RL. 1987. Endo-, para-, and ecodormancy: Physiological terminology and classification for dormancy research. HortScience: a publication of the American Society for Horticultural Science 22, 371–377.

Lavee S, May P. 1997. Dormancy of grapevine buds - facts and speculation. Australian journal of grape and wine research 3, 31–46.

van Leeuwen C, Schultz HR, Garcia de Cortazar-Atauri I, et al. 2013. Why climate change will not dramatically decrease viticultural suitability in main wine-producing areas by 2050. Proceedings of the National Academy of Sciences of the United States of America 110, E3051–2.

Londo JP, Johnson LM. 2014. Variation in the chilling requirement and budburst rate of wild Vitis species. Environmental and experimental botany 106, 138–147.

Londo JP, Kovaleski AP. 2017. Characterization of Wild North American Grapevine Cold Hardiness Using Differential Thermal Analysis. American journal of enology and viticulture, ajev.2016.16090.

Londo JP, Moyer MM, Mireles M, Mills L, Keller M, Workmaster BA, Atucha A, Kovaleski AP. 2023. Evaluation of Sample Preparation Practices Common with Differential Thermal Analysis of Grapevine Bud Cold Hardiness. American journal of enology and viticulture 74.

Luedeling E. 2012. Climate change impacts on winter chill for temperate fruit and nut production: A review. Scientia horticulturae 144, 218–229.

Luedeling E, Brown PH. 2011. A global analysis of the comparability of winter chill models for fruit and nut trees. International journal of biometeorology 55, 411–421.

Luedeling E, Kunz A, Blanke MM. 2013. Identification of chilling and heat requirements of cherry trees--a statistical approach. International journal of biometeorology 57, 679–689.

Marvel K, Su W, Delgado R, et al. 2023. Climate trends. Fifth National Climate Assessment November. https://nca2023.globalchange.gov/downloads/NCA5_Ch2_Climate-Trends.pdf.

Mills LJ, Ferguson JC, Keller M. 2006. Cold-Hardiness Evaluation of Grapevine Buds and Cane Tissues. American journal of enology and viticulture 57, 194–200.

Mo Y, Chen S, Wu Z, Tang J, Fu Y. 2024. The advancement in spring vegetation phenology in the northern hemisphere will reverse after 2060 under future moderate warming scenarios. Earth’s future 12, e2023EF003788.

Morales-Castilla I, García de Cortázar-Atauri I, Cook BI, Lacombe T, Parker A, van Leeuwen C, Nicholas KA, Wolkovich EM. 2020. Diversity buffers winegrowing regions from climate change losses. Proceedings of the National Academy of Sciences of the United States of America 117, 2864–2869.

Mozell MR, Thach L. 2014. The impact of climate change on the global wine industry: Challenges & solutions. Wine Economics and Policy 3, 81–89.

Nendel C. 2010. Grapevine bud break prediction for cool winter climates. International journal of biometeorology 54, 231–241.

North M, Workmaster BA, Atucha A. 2021. Cold Hardiness of Cold Climate Interspecific Hybrid Grapevines Grown in a Cold Climate Region. American journal of enology and viticulture 72, 318–327.

North M, Workmaster BA, Atucha A. 2022. Effects of chill unit accumulation and temperature on woody plant deacclimation kinetics. Physiologia plantarum 174, e13717.

Pagter M, Arora R. 2013. Winter survival and deacclimation of perennials under warming climate: physiological perspectives. Physiologia plantarum 147, 75–87.

Parker A, de Cortázar-Atauri IG, Chuine I, et al. 2013. Classification of varieties for their timing of flowering and veraison using a modelling approach: A case study for the grapevine species Vitis vinifera L. Agricultural and Forest Meteorology 180, 249–264.

Parker AK, de Cortázar-Atauri IG, Van Leeuwen C, Chuine I. 2011. General phenological model to characterise the timing of flowering and veraison of Vitis vinifera L. Australian journal of grape and wine research 17, 206–216.

Pellegrino A, Blackmore D, Clingeleffer P, Walker R. 2022. Comparison of methods for determining budburst date in grapevine. OENO One 56, 73–86.

Pellegrino A, Rogiers S, Deloire A. 2020. Grapevine latent bud dormancy and shoot development. IVES Technical Reviews, vine and wine doi: 10.20870/ives-tr.2020.3420.

Pertille RH, Citadin I, Patto LS, Oldoni TLC, Scariotto S, Grigolo CR, Lauri P-É. 2021. High-chilling requirement apple cultivar has more accentuated acrotony than low-chilling one in mild winter region. Trees 35, 1135–1150.

Piña-Rey A, Ribeiro H, Fernández-González M, Abreu I, Rodríguez-Rajo FJ. 2021. Phenological model to predict budbreak and flowering dates of four Vitis vinifera L. cultivars cultivated in DO. Ribeiro (north-west Spain). Plants 10, 502.

Rahemi A, Fisher H, Dale A, Taghavi T, Kelly J. 2021. Bud dormancy pattern, chilling requirement, and cold hardiness in*Vitis vinifera*L. ‘Chardonnay’ and ‘Riesling’. Canadian journal of plant science. Revue canadienne de phytotechnie 101, 871–885.

Reynolds AG, Reisch BI. 2015. Grapevine breeding in the Eastern United States. Grapevine Breeding Programs for the Wine Industry. Elsevier, 345–358.

Ritz C, Baty F, Streibig JC, Gerhard D. 2015. Dose-response analysis using R. PloS one 10, e0146021.

Rochette P, Bélanger G, Castonguay Y, Bootsma A, Mongrain D. 2004. Climate change and winter damage to fruit trees in eastern Canada. Canadian journal of plant science. Revue canadienne de phytotechnie 84, 1113–1125.

Rubio S, Dantas D, Bressan-Smith R, Pérez FJ. 2016. Relationship Between Endodormancy and Cold Hardiness in Grapevine Buds. Journal of plant growth regulation 35, 266–275.

Sandor ME, Aslan CE, Pejchar L, Bronstein JL. 2021. A mechanistic framework for understanding the effects of climate change on the link between flowering and fruiting phenology. Frontiers in ecology and evolution 9, 752110.

Segrestin J, Bernard-Verdier M, Violle C, Richarte J, Navas M-L, Garnier E. 2018. When is the best time to flower and disperse? A comparative analysis of plant reproductive phenology in the Mediterranean. Functional ecology 32, 1770–1783.

Vitasse Y, Lenz A, Körner C. 2014. The interaction between freezing tolerance and phenology in temperate deciduous trees. Frontiers in plant science 5, 541.

Vyse K, Pagter M, Zuther E, Hincha DK. 2019. Deacclimation after cold acclimation-a crucial, but widely neglected part of plant winter survival. Journal of experimental botany 70, 4595–4604.

Wang X, Li H, García de Cortázar Atauri I. 2020. Assessing grapevine phenological models under Chinese climatic conditions. OENO One doi: 10.20870/oeno-one.2020.54.3.3195.

Wang H, Moghe GD, Kovaleski AP, et al. 2023. NYUS.2: an Automated Machine Learning Prediction Model for the Large-scale Real-time Simulation of Grapevine Freezing Tolerance in North America. bioRxiv, 2023.08.21.553868.

Wisniewski M, Bassett C, Gusta LV. 2003. An overview of cold hardiness in woody plants: Seeing the Forest through the trees. HortScience: a publication of the American Society for Horticultural Science 38, 952–959.

Wisniewski M, Nassuth A, Teulières C, Marque C, Rowland J, Cao PB, Brown A. 2014. Genomics of cold hardiness in woody plants. Critical reviews in plant sciences 33, 92–124.

Wolf TK, Cook MK. 1994. Cold hardiness of dormant buds of grape cultivars: Comparison of thermal analysis and field survival. HortScience 29, 1453–1455.

Yilmaz T, Alahakoon D, Fennell A. 2020. Freezing tolerance and chilling fulfillment differences in cold climate grape cultivars. Horticulturae 7, 4.

